# Functional effects of variation in transcription factor binding highlight long-range gene regulation by epromoters

**DOI:** 10.1101/620062

**Authors:** Joanna Mitchelmore, Nastasiya Grinberg, Chris Wallace, Mikhail Spivakov

## Abstract

Identifying DNA cis-regulatory modules (CRMs) that control the expression of specific genes is crucial for deciphering the logic of transcriptional control. Natural genetic variation can point to the possible gene regulatory function of specific sequences through their allelic associations with gene expression. However, comprehensive identification of causal regulatory sequences in brute-force association testing without incorporating prior knowledge is challenging due to limited statistical power and effects of linkage disequilibrium. Sequence variants affecting transcription factor (TF) binding at CRMs have a strong potential to influence gene regulatory function, which provides a motivation for prioritising such variants in association testing. Here, we generate an atlas of CRMs showing predicted allelic variation in TF binding affinity in human lymphoblastoid cell lines (LCLs) and test their association with the expression of their putative target genes inferred from Promoter Capture Hi-C and immediate linear proximity. We reveal over 1300 CRM TF-binding variants associated with target gene expression, the majority of them undetected with standard association testing. A large proportion of CRMs showing associations with the expression of genes they contact in 3D localise to the promoter regions of other genes, supporting the notion of ‘epromoters’: dual-action CRMs with promoter and distal enhancer activity.

## Introduction

Identifying DNA cis-regulatory modules (CRMs) that control the expression of specific genes is crucial for deciphering the logic of transcriptional control and its aberrations. Advances of the last decade have made it possible to predict active CRMs based on chromatin features (1, 2) and detect the binding of dozens of TFs to these regions (3, 4). However, deletion of known or predicted CRMs often shows no observable phenotype, suggesting that some CRMs either lack appreciable gene regulatory function or are efficiently buffered by other sequences, at least under normal conditions (5–9). In addition, the sequence, chromatin state and genomic location of CRMs do not immediately provide information on their target genes (10). Therefore, evidence from complementary approaches is required to establish the function of specific CRMs in transcriptional control.

Natural genetic variation can theoretically provide a direct indication of gene regulatory function by revealing the allelic associations between specific variants and gene expression (11, 12). While expression quantitative trait loci (eQTLs) identified this way have provided important insights into gene control and the mechanisms of specific diseases (13, 14), a number of challenges hamper comprehensive detection of functional sequences in ‘brute-force’ eQTL testing (15, 16). In particular, the immense search space leads to a heavy multiple testing burden resulting in reduced sensitivity. This problem is typically mitigated in part by testing for ‘cis-eQTLs’ separately within a limited distance window (~100kb); this distance range is, however, an order of magnitude shorter than that of known distal CRM activity (17–19). In addition, correlation structure arising from linkage disequilibrium (LD) requires disentangling causal from spurious associations, which is particularly challenging in the likely scenario when multiple functional variants with modest effects co-exist within the same LD block (20). These challenges provide a strong motivation for incorporating prior knowledge into association testing for identifying causal regulatory variants.

The recruitment of transcription factors (TFs) to CRMs plays a key role in their regulatory function (21, 22), and mutations leading to perturbed TF binding are known to underpin developmental abnormalities and disease susceptibility (18, 23, 24). Therefore, sequence variation affecting TF binding affinity at CRMs has a strong potential to have causal influence on their regulatory function and can therefore provide insights into the logic of gene control. Variation in TF binding across multiple individuals has been assessed directly for several TFs (25–30), but high resource requirements of these analyses limit the number of TFs and individuals profiled this way. Alternatively, the effects of local sequence variation on TF binding can be predicted, at least in part, based on prior information regarding the TFs’ DNA binding preferences. The representation of such preferences in the form of position weight matrices (PWMs) (31) has proven particularly useful, as it provides a quantitative measure of how much a given sequence substitution is likely to perturb TF binding consensus. Consistent with this, we and others have previously shown that the specificity of TF binding preferences to a given motif position correlates with the functional constraint of the underlying DNA sequences, both within and across species (32–34). Classic PWM-based approaches to TF binding prediction focused on identifying short sequences showing a non-random fit to the PWM model compared with background (35, 36). More recently, biophysical modelling of TF binding affinity (37, 38) has provided a natural framework to extend this analysis by integrating over all PWM match signals within a DNA region (39, 40), including those from lower-affinity sites that are a known feature of many functional CRMs (41–43).

Long-range CRMs such as gene enhancers commonly act on their target promoters through DNA looping interactions (44, 45). Therefore, information on three-dimensional chromosomal organisation enables predicting the putative target genes of these elements (46, 47) and thus has the potential to significantly improve the functional interpretation of regulatory variation. Approaches that couple chromosome conformation capture with target sequence enrichment such as Promoter Capture Hi-C (48–50) are particularly useful in this regard, as they make it possible to detect regulatory interactions globally and at high resolution with reasonable amounts of sequencing (51–59).

Here we integrate TF binding profiles in a human lymphoblastoid cell line (LCL) (4) with patterns of natural sequence variation (60) to generate an atlas of CRMs predicted to show significant TF binding variability across LCLs derived from multiple individuals. We delineate the putative target genes of these CRMs from their interactions with gene promoters based on Promoter Capture Hi-C and linear proximity (49, 61), and test for associations between the CRMs’ TF-binding affinity and target gene expression using transcriptomics data for hundreds of LCLs (62). Prioritising CRMs that show predicted variation in TF-binding affinity based on a biophysical model (39, 40) makes it feasible to perform association analysis in a manner that accounts for multiple variants affecting the binding of the same TF, as well as for multiple CRMs targeting the same gene. Using this approach, we reveal over 1300 CRM variants associated with expression of specific genes, the majority of them undetected with conventional eQTL testing at a standard FDR threshold. We find that a large proportion of CRMs showing associations with the expression of distal genes localise in the immediate vicinity of the TSSs of other genes and connect to their targets via DNA looping interactions, suggesting their role as ‘epromoters’: the recently identified dual-action regulatory regions with promoter and distal enhancer activity (63–65).

## Materials and Methods

### CRM definition

ChIP-seq narrow peak files for 52 TFs in GM12878 were downloaded from the UCSC ENCODE portal (4). Where multiple datasets were available for the same TF, the intersect of the ChIP-seq peaks was taken for all TFs except ERG1, for which we took the union of the two datasets available, since one of them had substantially fewer peaks than the other. CRMs were defined by taking the union of the peaks for the 52 TFs with a minimum overlap of one base pair.

### Detection of TF binding affinity variants

Variant calls for 359 LCLs of European ancestry (CEU, TSI, FIN, GBR, and IBS) that overlapped with the CRMs defined as above were downloaded from the 1000 Genomes Project (release Phase 3; 20130502) (60). Multi-allelic variants and variants with a minor allele frequency < 5% were removed. Unique haplotypes (i.e., unique combinations of SNPs/indels) were identified across the 359 LCLs individuals for each CRM. The GRCh37 genomic sequence for each CRM (accessed using the Bioconductor package BSGenome (https://doi.org/10.18129/B9.bioc.BSgenome) was then patched to create the sequence for each unique haplotype.

For each TF detected as bound at a given CRM in GM12878 (based on ChIP-seq data), we computed the affinity for each haplotype and each PWM for this TF available from ENCODE (66) using the TRAP biophysical model (39), as implemented in the R package tRap (https://github.com/matthuska/tRap). Default parameters were used, with the exception of setting pseudocount to zero, since we were using frequency as opposed to count matrices. We chose TRAP over a motif hit-based approach, as it naturally incorporates the effects of multiple low affinity sites and multiple variants per CRM.

CRM binding affinities were normalised using a method proposed by Manke et al. (40), such that changes in them could be compared between different PWMs. Briefly, CRM affinities are converted to statistical scores (*A*) representing the probability of observing a given or higher affinity for a given TF in the background sequence (note that lower values of *A* therefore reflect higher affinities). Binding affinities are parameterised using the extreme value distribution whose parameters are estimated for a range of background sequences encompassing the lengths of all CRMs (40, 100, 200, 250, 300, 400, 500, 800, 1000, 2000, 3000) using the fit.gev function in the the tRap R package. CRMs not bound by a given TF are cut/extended to the required length and used as background sequences.

For all CRM-TF/PWM combinations with *A*<0.1 in the highest-affinity allele of GM12878, we computed the log-fold change in affinity between all observed haplotypes and the highest-affinity allele of GM12878 for the given PWM:

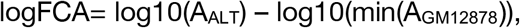

where min(*A*_GM12878_) is the normalised affinity of the highest-affinity allele in GM12878 cells, and *A*_ALT_ is the normalised affinity of the alternative haplotype. For instances where *A*_ALT_ or *A*_GM12878_ for a given PWM was zero, the lowest observed non-zero normalised affinity for that PWM across all CRMs was used instead. The logFCAs for multiple PWMs of the same TF were then combined by taking the median. Overall, this approach produced a single logFCA for each TF binding affinity haplotype at each CRM. We shall refer to this quantity as the “log-ratio” in the Results section.

### DNase I sensitivity QTL (dsQTL) analysis

The dsQTL dataset from (67) lists significant associations between normalized DNase-seq read depth (binned in 100bp non-overlapping windows) and the genotypes of SNPs/indels within 1kb of the DHS in 70 Yoruban LCLs. We downloaded this dataset from Gene Expression Omnibus (accession number GSE31388), and converted it to GRCh37 using liftOver (68). For all CRMs with a predicted logFCA > 0 for at least one TF, the individual effect of all SNPs at the CRM on TF affinity was calculated. CRMs were then filtered for those, where the SNP causing the largest change in TF affinity (“driver SNP”) had a MAF<0.05 in the 70 individuals from (67). We then counted the number of overlaps between these CRMs and the 100bp DNase HS windows (minimum overlap 1bp), repeating this for CRMs filtered according to successively larger logFCA thresholds. To estimate expected overlap, for each threshold, we randomly sampled a control set of CRMs 1000 times, matching the sample size and “driver” SNP allele frequency distribution to the test set at a given threshold, and overlapped this set with DNase HS windows in the same way as the test set.

### Linking of CRMs with target genes

Promoter Capture Hi-C data for GM12878 were obtained from Mifsud et al. (49). Significant interactions were re-called at a *HindIII* restriction fragment level using the CHiCAGO pipeline (61), with a CHiCAGO score cutoff of five (CHiCAGO scores correspond to soft-thresholded, −log weighted p-values against the background model). Baits were annotated for transcriptional start sites (TSSs) using the bioMart package in R (69) based on Ensembl TSS data for GRCh37 reference assembly. Baits containing TSSs for more than one gene were excluded (4,178 out of 22,076), leaving 17,898 baits in the analysis. CRMs were assigned to target promoters by overlapping with the promoter-interacting regions (PIRs) of significant interactions (“distal” CRMs). Restriction fragments immediately flanking the promoter fragment are excluded from Promoter Capture Hi-C analysis, creating a “blind window”. Therefore, we additionally called “proximal” CRMs using a window-based approach, assigning all CRMs located within within 9kb of the midpoint of the promoter-containing fragment to the respective promoter.

### Gene expression data processing

We downloaded PEER-normalised (70) gene-level RPKMs for 359 EUR LCLs profiled in the GEUVADIS project (62) from ArrayExpress (71) (accession E-GEUV-3). The data were filtered to expressed genes by removing genes with zero read counts in >50% of samples. For expression association testing by linear regression, the PEER-normalised residuals for each gene were further rank-transformed to standard normal distribution, using the rntransform function in the R package GenABEL (72).

### Association between TF binding affinity variants and gene expression: thresholded approach

In this approach, we classified each predicted TF-binding affinity CRM haplotype as either “high” or “low” affinity based on a threshold. In some instances, however, using a hard threshold to classify alleles can result in alleles with very similar log-fold affinity changes being differentially classified, which can obscure true affinity-expression associations. To avoid this, we used a dynamic thresholding approach, where for each affinity variant we set the threshold logFCA_0_ to 80% of the value of the 85^th^ percentile of variants less than or equal to the hard threshold of −0.3. All alleles with logFCA <= logFCA_0_ were taken as low affinity. Alleles with either logFCA > logFCA_0_/4 (for logFCA_0_/4 > −0.3) or logFCA > −0.3 were taken as high affinity. Note this resulted in some alleles classified as neither high nor low affinity. Individuals containing at least one unclassified allele for a given TF/CRM were excluded from the testing for the respective association (the number of individuals tested for each association is listed in Table S1).

A regression model was then fitted using TF-binding affinity CRM haplotypes as predictors of the expression level of their target genes (presented in terms of normalised PEER residuals). Suppose that a gene is targeted by *K* predicted TF-affinity CRM variants, denoted as *X* = (*X*_1_, *X*_2_,…, *X*_*K*_,), which are encoded as the number of copies of the low affinity allele carried by each individual. The regression model is fitted as follows:

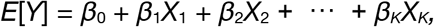

where *E*[*Y*] is the expected value of the normalised PEER residuals *Y*. Where multiple predicted TF affinity CRM variants targeting a given gene were in perfect correlation (|*β*|>0.99), they were collapsed into a single predictor.

ANOVA was used to test the overall significance of each regression model, with multiple testing correction performed on the gene-level p-values by FDR estimation. For genes showing significant associations in models with multiple TF binding affinity variants as predictors, t-tests were performed to identify variants with regression coefficients significantly different from zero. Variants with unadjusted coefficient-level p-values<0.05 were taken to be significantly associated with target gene expression, conditional on significant gene-level association at 10% FDR.

### Association between TF binding affinity variants and gene expression: threshold-free approach

In this approach, we performed multiple regression using PEER expression residuals for each gene as the response variable, this time using the sum of logFCA across both alleles for each individual for each TF affinity CRM variant as predictors instead of thresholded CRM haplotypes. For each gene, all distal and proximal CRMs with logFCA > 0 were included. As with the thresholded approach, ANOVA was used to test the significance of each gene model, and genes showing associations at 10% FDR were considered significant.

Due to high collinearity among the predicted affinity changes, to identify specific CRM variants significantly associated with target gene expression we used elastic net regression for each significantly associated gene (λ_2_=0.5). The significance of each predictor as it entered the model was then tested using a method by Lockhart et al. (73) and implemented in the covTest R package (https://cran.r-project.org/src/contrib/Archive/covTest/covTest_1.02.tar.gz). Variants that entered the model with p<0.05 and remained in the model were taken as significant.

### eQTL fine-mapping

We fine-mapped eQTL causal variants in the LCL expression data within a window of +/−200kb of each CRM using a Bayesian stochastic search method that allows for multiple causal variants, GUESSFM (https://github.com/chr1swallace/GUESSFM) (74). This requires a prior on the number of causal variants per region, which we set as Bin(*n*,2/*n*) where *n* is the number of variants in the fine mapping window. This setting gives a prior expectation of 2 causal variants per region but allows all values from 0 to *n*. We visually checked traces to ensure the MCMC samples had converged. Raw GUESSFM data have been uploaded to OSF (https://osf.io/e5vsh/).

To estimate the proportion of possibly causal eQTLs identified by GUESSFM (marginal posterior probability of inclusion [mppi]>> 0.001) among the TF-binding affinity variants showing the strongest eQTL signal per CRM (“test SNPs”), we compared it with the same proportion obtained for “random SNPs”. The “random SNPs” were sampled from the same +/−200kb windows around CRMs, matching the distribution of their minor allele frequencies to that across the “test SNPs”.

### Causal variant colocolisation analysis

An association between an epromoter variant and the expression of both a proximal and a distal gene may indicate that this variant is causal for the expression of both genes. However, the same association may arise from distinct causal variants for each gene that are in LD with each other and are tagged by the same epromoter variant. To differentiate between these situations, we used the Bayesian colocalisation technique coloc (75). Coloc evaluates the posterior probabilities of five mutually exclusive hypotheses: no association of any variant in the region with either trait (H0), association with first trait but not the second (H1), association with second trait but not the first (H2), two separate causal variants (H3), and, finally, a unique shared causal variant (H4). Coloc assumes at most one casual variant per locus. To mitigate this limitation, where there was evidence for multiple causal variants, we tested for colocalisation between all pairs of signals for each gene, by conditioning out the other signals. Coloc has also been originally designed for testing two sets of associations measured on different individuals. Therefore, before running it on the data measured in the same individuals (i.e., the expression of the proximal and distal gene across the 359 CEU LCLs) we confirmed by simulation that for a quantitative trait the results appear robust to correlated errors (Figure S1).

## RESULTS

### An atlas of CRMs with predicted variation in TF binding affinity in LCLs

We used the ChIP-seq binding profiles of 52 TFs profiled by the ENCODE project (4) in GM12878 LCL to define 128,766 CRMs in these cells, merging across overlapping ChIP regions for multiple TFs (Figure 1). Just over half (55%) of CRMs defined this way were bound by more than a single TF. For 41/52 TFs with known PWMs, we then used a biophysical model (39) to estimate their binding affinity to each allele of each CRM in GM12878, pooling information across multiple PWMs for the same TF where available (see Materials and Methods). To enable the comparison of binding affinities between different TFs, we expressed them relative to the respective ‘background’ affinities using an approach based on the generalised extreme value distribution (40) (see Materials and Methods for details).

**Figure 1.**
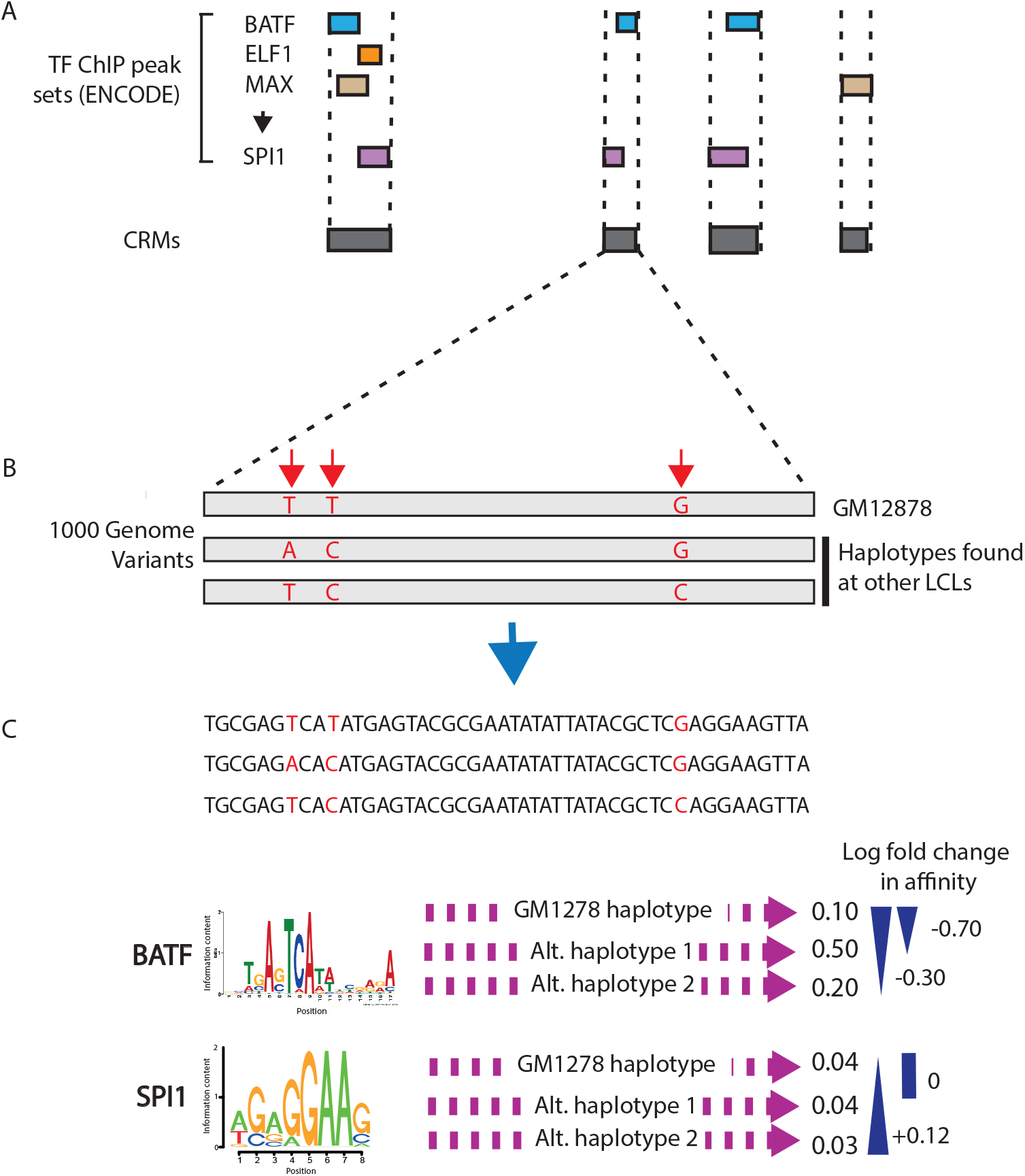
Definition of TF binding affinity variants. (A) TF ChIP-Seq data for 52 TFs profiled by the ENCODE project (4) in GM12878 are used to define CRMs, merging across overlapping ChIP regions for multiple TFs. Here ChIP seq regions for four TFs are depicted by different coloured rectangles. (B) Using variants from the 1000 Genomes Project, unique haplotypes for each CRM across the 359 LCLs are identified, including the haplotype/s of GM12878. (C) For each TF bound at the CRM in GM12878, ENCODE PWMs for the given TF are used to predict the normalised binding affinity for each unique haplotype (note that lower affinity values reflect higher affinities). The log fold affinity change between the highest affinity haplotype of GM12878 and each alternative haplotype is computed for each PWM and for each haplotype/TF the median log fold affinity change across all PWMs belonging to the given TF is taken (here only one PWM per TF is depicted).

We next asked how natural genetic variation at CRMs affects their TF-binding affinity. For this, we took advantage of the genotypes of an additional 358 LCLs also derived from European-ancestry individuals that are available from the 1000 Genomes project (60). We then calculated a TF affinity log-ratio between each alternative haplotype and the highest-affinity haplotype of GM12878 (Figure 1; see Materials and Methods). Overall, 38,804 CRMs had one or more alternative haplotypes with predicted changes in binding affinity for at least one TFs (affinity log-ratios ranging between −12.9 and 13.17). We have made the full atlas of TF-binding CRM variants publicly available at https://osf.io/fa4u7.

### CRMs showing TF-binding variation are enriched for chromatin accessibility variants

TF binding is known to associate with increased chromatin accessibility. Therefore, to validate our predicted changes in TF-binding affinity, we took advantage of a published study (67) that profiled chromatin accessibility across 70 LCLs using DNase-seq and identified ~9,000 significant associations between DNase-seq signal and genotype (“DNase I sensitivity QTLs”, dsQTLs). If our predicted TF affinity variants reflected real changes in TF binding affinity, we would expect them to show enrichment at regions of differential chromatin accessibility. To verify this, we quantified enrichment of differential chromatin accessibility at sets of CRMs showing predicted TF affinity variation above successively larger thresholds. As can be seen from Figure 2, CRMs with non-zero differences in TF-binding affinity across LCLs showed a significant enrichment at differential DNAse I sensitivity regions compared with a matched random set of CRMs (permutation test p<0.001, see Materials and Methods for details). Moreover, this enrichment increased with the magnitude of the predicted affinity change (Figure 2). These results provide direct functional evidence that our approach adequately predicts changes in TF binding associated with genetic variation. It should be noted that the observed overlap in absolute terms is likely underestimated, due to the relatively limited DNase-seq sequencing depth and sample size in the available dataset (67).

**Figure 2.**
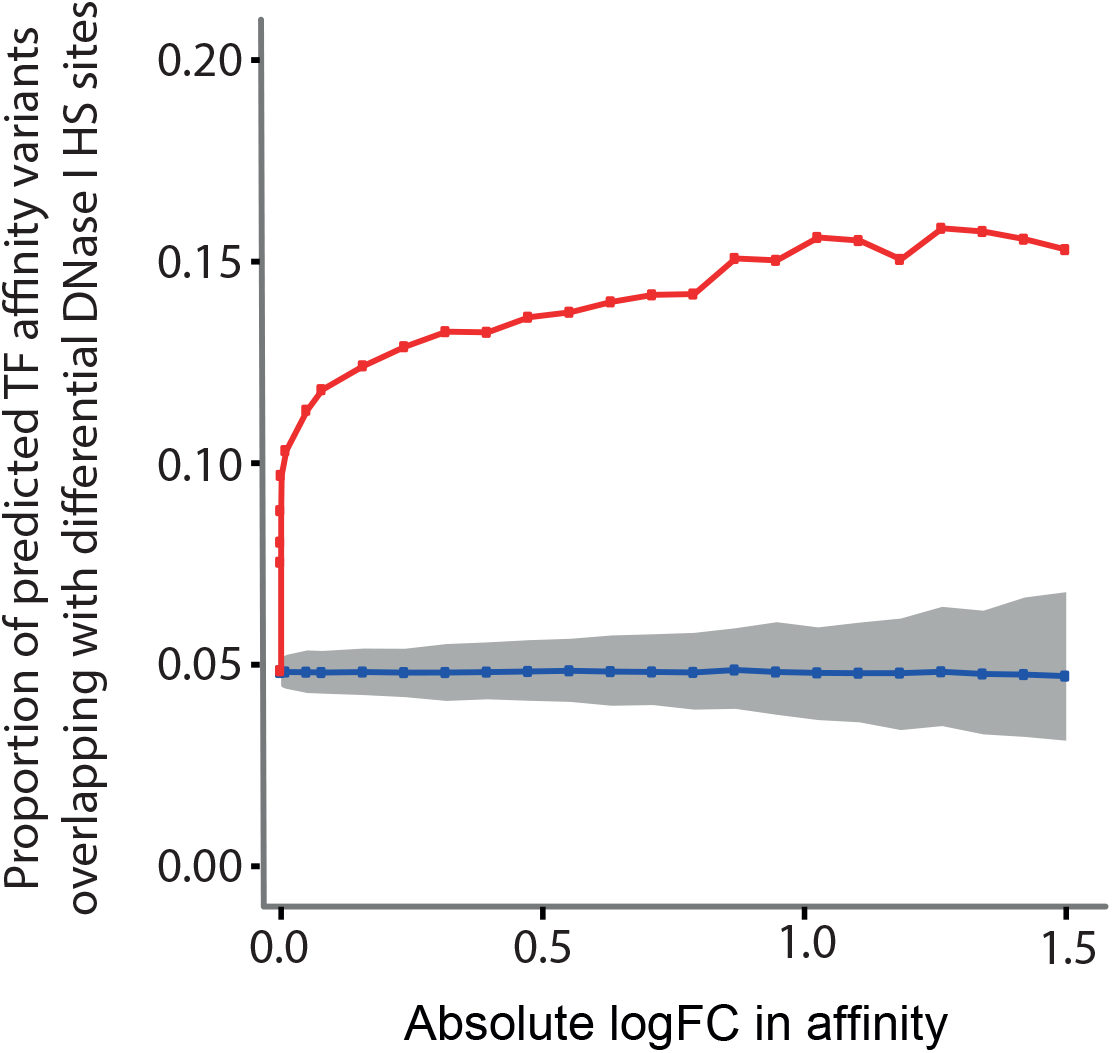
TF binding affinity variants are enriched at regions showing variation in DNase I hypersensitivity. The proportion of CRMs with log fold affinity changes over a range of thresholds that overlap with differential DNase I hypersensitivity (HS) sites (identified as dsQTLs in ref. (67)) are depicted by red squares. CRMs were filtered to those where the SNP driving the affinity change has a MAF > 5% in the 70 YRI individuals. The mean proportion of randomly sampled CRMs that overlap with differential DNase I HS sites across 1000 permutations are shown in blue, with the grey ribbon showing the range where 90% of the permuted values (number of overlaps) lie. For each threshold the control sets of CRMs were matched in sample size and “driving” SNP allele frequency distribution to the number and allele frequency distribution respectively of the predicted affinity variants over the corresponding threshold.

### Variation in TF binding affinity at CRMs associates with target gene expression

To identify quantitative associations between TF binding variation at CRMs and the expression of their target genes, we used genome-wide gene expression data from the GEUVADIS project (62) that included 358/359 of the LCLs used in our analysis (with the exception of GM12878). In contrast to traditional eQTL testing, here we devised an approach that prioritises TF-binding variants and their putative target genes *a priori* and performs testing at the CRM level. In total, we selected 3,285 CRMs with predicted variation in the binding for at least one TF (log-ratio > 0.3). We then tested the association of each CRM haplotype with the expression levels of their target genes defined on the basis of 3D interactions or close spatial proximity (within 9kb; see Materials and Methods). As evidence of 3D promoter-CRM interactions, we used high-resolution Promoter Capture Hi-C (PCHi-C) data in GM12878 cells (49, 61). The highly reduced search space has enabled testing for associations at the gene level, with all CRMs targeting the same gene and showing TF-binding variation included into the regression model (see Materials and Methods). This approach identified 245 “eGenes” with significant associations between predicted TF-binding affinity at CRMs and gene expression (16% of 1530 genes tested, at 10% FDR; Table S1). In total, 161 “proximal” (within 9 kb) and 101 “distal” TF-CRM affinity variants (with contacts detected by PCHi-C) were found to underlie these associations, corresponding to 26% and 6% of all variants tested, respectively (t-test p-value <0.05; Table S1). Figure 3 shows an example of the detected association between the expression of *KLF6* and variation in the binding affinity of BATF transcription factor at a distal CRM that is located 88 kb away from *KLF6* promoter and contacts it in 3D according to PCHi-C (gene-level FDR=1.21×10^−2^, BATF variant p-value=5.16×10^−4^, effect size=0.26). Individuals homozygous for the high-affinity BATF binding allele showed the lowest levels of *KLF6* expression, while those homozygous for the low-affinity BATF binding alleles showed the highest levels (Figure 3). This suggests that BATF acts as a negative regulator of *KLF6* expression, consistent with its known role as a repressor of AP-1-dependent transcriptional activity (76).

**Figure 3.**
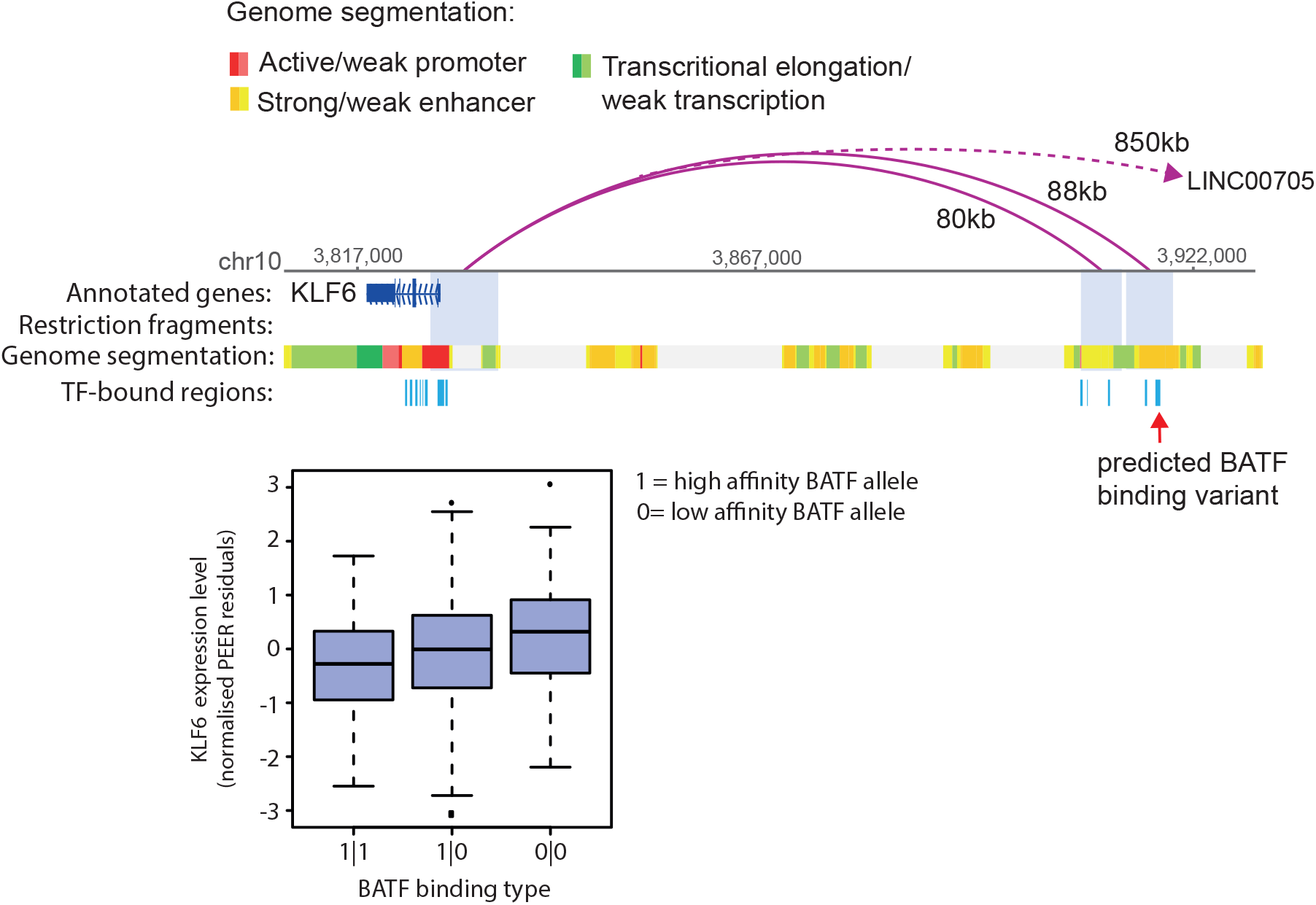
Example of association between a TF-binding affinity CRM variant and gene expression. (A) Genome browser representation of the distal interactions (pink arches) of *KLF6* promoter in the LCL GM12878, as detected by Promoter Capture Hi-C (49). Two out of the three fragments interacting with KLF6 are shown; the third fragment, which is located 850kb away from the KLF6 promoter and contains the gene LINC00705, was omitted due to space constraints. Genome segmentation tracks for GM12878 are shown (108). CRMs at the two distally interacting fragments and TSS-proximal window are depicted in azure blue. The far-right CRM, which interacts with the KLF6 promoter 88kb away, is predicted to impact BATF binding affinity across the 359 LCLs. (B). Boxplot showing the association in LCLs between mRNA levels (as measured with RNA-seq by the GEUVADIS consortium) and predicted BATF affinity CRM haplotype. KLF6 expression is significantly associated with BATF binding type (gene level FDR adjusted p-value=1.21×10^−2^, BATF variant p-value= 5.16×10^−4^, effect size=0.26).

A total of 420/1530 genes (27%) were linked with multiple predicted TF-binding variants (either for different TFs bound at the same CRM or at different CRMs). For 16 of these genes, we detected significant associations between more than one such variant and the expression level. One example is the nuclear receptor gene *NR2F6* whose expression significantly associated with predicted variation in the binding affinities of SMC3 and SRF to distal CRMs located, respectively, 41 kb and 19 kb away (Figure 4; gene-level FDR = 4.06×10^−7^, SMC3 effect size=0.26, p-value=3×10^−4^; SRF effect size=0.61, p-value=1.19×10^−7^).

**Figure 4.**
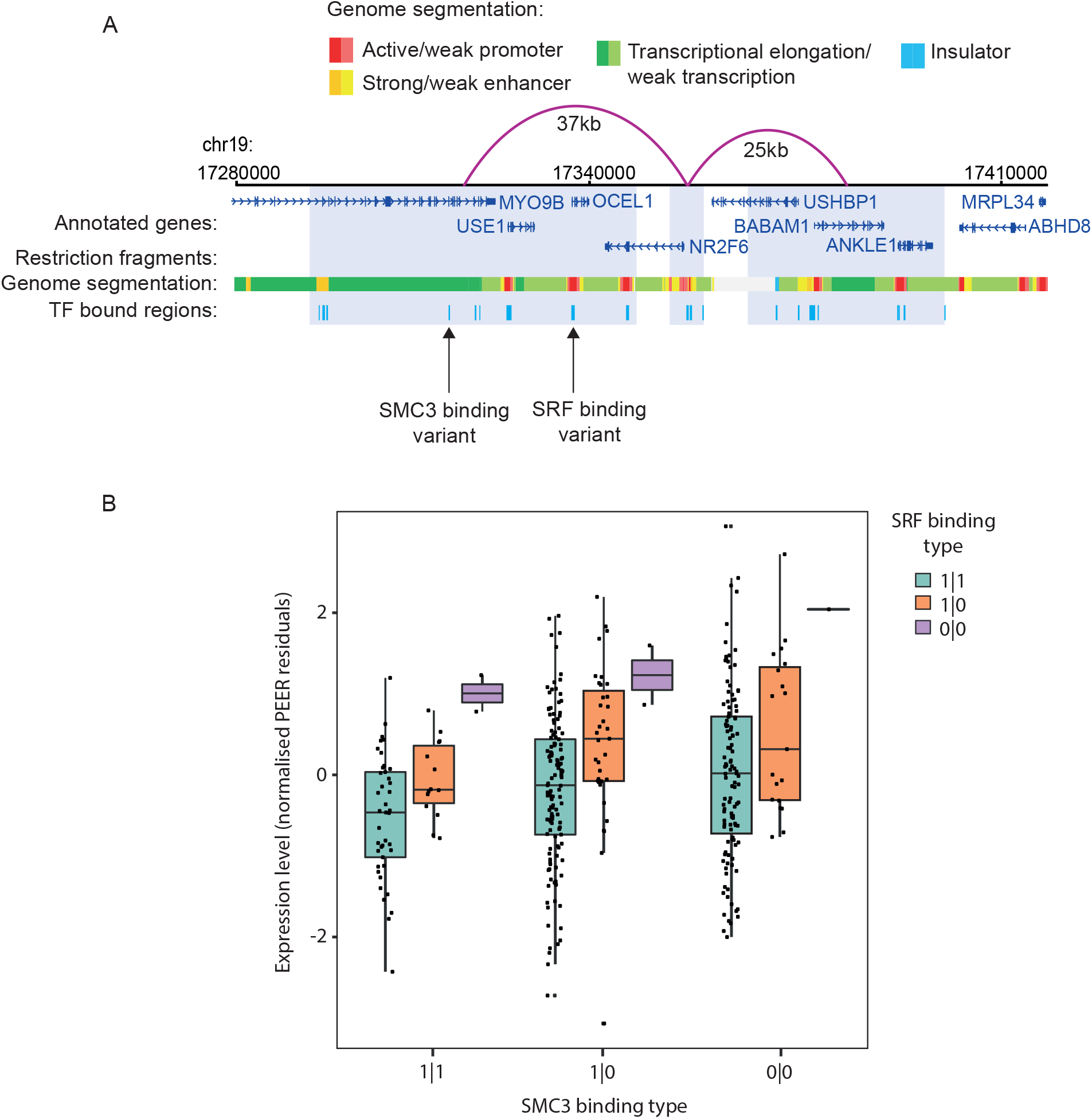
Example of a multi-variant association between TF binding affinity at CRMs and their target gene expression. (A) Genome browser representation of NR2F6 promoter distal interactions (represented by pink arches) as detected by promoter capture Hi-C (49) in LCL GM12878. The genome segmentation track for GM12878 based on chromHMM (108) is also shown. CRMs at the distally interacting fragments (pale blue) and NR2F6 TSS-proximal window are depicted in azure blue. The distal fragment downstream of NR2F6 contains two predicted affinity variant CRMs: one 44kb away from the NR2F6 promoter and the other 19kb away, predicted to impact SMC3 and SRF binding affinity respectively across the 359 LCLs. (B) Association between NR2F6 mRNA levels and predicted SMC3 and SRF affinity haplotypes.

Owing to the *a priori* prioritisation of variants for association testing in our approach (i.e., testing only variants predicted to impact TF binding), we carried out far fewer association tests than in a standard eQTL analysis, thus reducing the multiple testing burden and increasing sensitivity. We therefore asked if we were able to detect additional associations compared with those reported for a standard eQTL analysis performed by the GEUVADIS project (note that this analysis also used an additional 103 LCLs not included in our study, which were either of non-European ancestry or not genotyped in 1000Genomes). To compare our CRM-based association results to GEUVADIS eQTL SNPs, we identified the SNP causing the largest change in affinity for the respective TF at each CRM (192 eQTL SNPs in total at 5% FDR to match the FDR level used by GEUVADIS). Of these, 78 SNPs (42%) were detected as significant by GEUVADIS. Therefore, the remaining 114/192 (58%) eQTL SNPs identified in our approach corresponded to not previously reported associations.

### Threshold-free testing based on TF-binding affinities reveals further expression associations

The analysis above was performed broadly within the conventional paradigm of eQTL testing, whereby expression was compared across three diploid genotypes (two homozygous and one heterozygous), except that these genotypes corresponded to cases whereby variation was predicted to appreciably disrupt TF binding based on a predefined threshold (we shall refer to this approach as “thresholded”). However, since TF-binding affinity haplotypes were defined at the CRM level, more than two alleles were commonly observed per CRM (in 12-100% cases depending on the TF). In the thresholded approach, we pooled multiple alleles into either “high-affinity” or “low-affinity” haplotypes and disregarded outliers (see Materials and Methods). We reasoned, however, that is it also possible to regress gene expression against normalised TF-binding affinities directly without thresholding and haplotype pooling, leading to increased precision and sensitivity of association testing. As expected, this threshold-free approach revealed a considerably larger number of genes significantly associated with CRM affinity variants (1033 at 10% FDR compared with 245 detected in the “thresholded” approach above).

One challenge arising in the threshold-free approach is that it leads to many more binding variants tested for each gene (both within the same CRM and across CRMs) that are often in linkage disequilibrium (LD) with each other, leading to collinearity in the regression models. Therefore, to detect significant associations at CRM level, we performed elastic net regression for each of the 895/1033 identified eGenes that were targeted by multiple CRMs with predicted TF binding variants. To ascertain the significance of regression coefficients in elastic net regression, we used a covariance test for adaptive linear models (73), identifying 1328 significant CRM-gene associations for the 895 eGenes tested (Table S2; see Materials and Methods for details). One example of a newly identified association is between a nucleotide transporter gene *SLC29A3* and the binding affinity of SIN3A at a CRM overlapping with the TSS of *SLC29A3* (gene-level FDR=1.60×10^−4^). Five alternative SIN3A binding affinity haplotypes were observed across the 358 LCLs, with log fold-changes in affinity for SIN3A ranging from −0.037 to 0.001 (elastic net effect size=-0.14, p-value ~ 0; Figure 5A).

**Figure 5.**
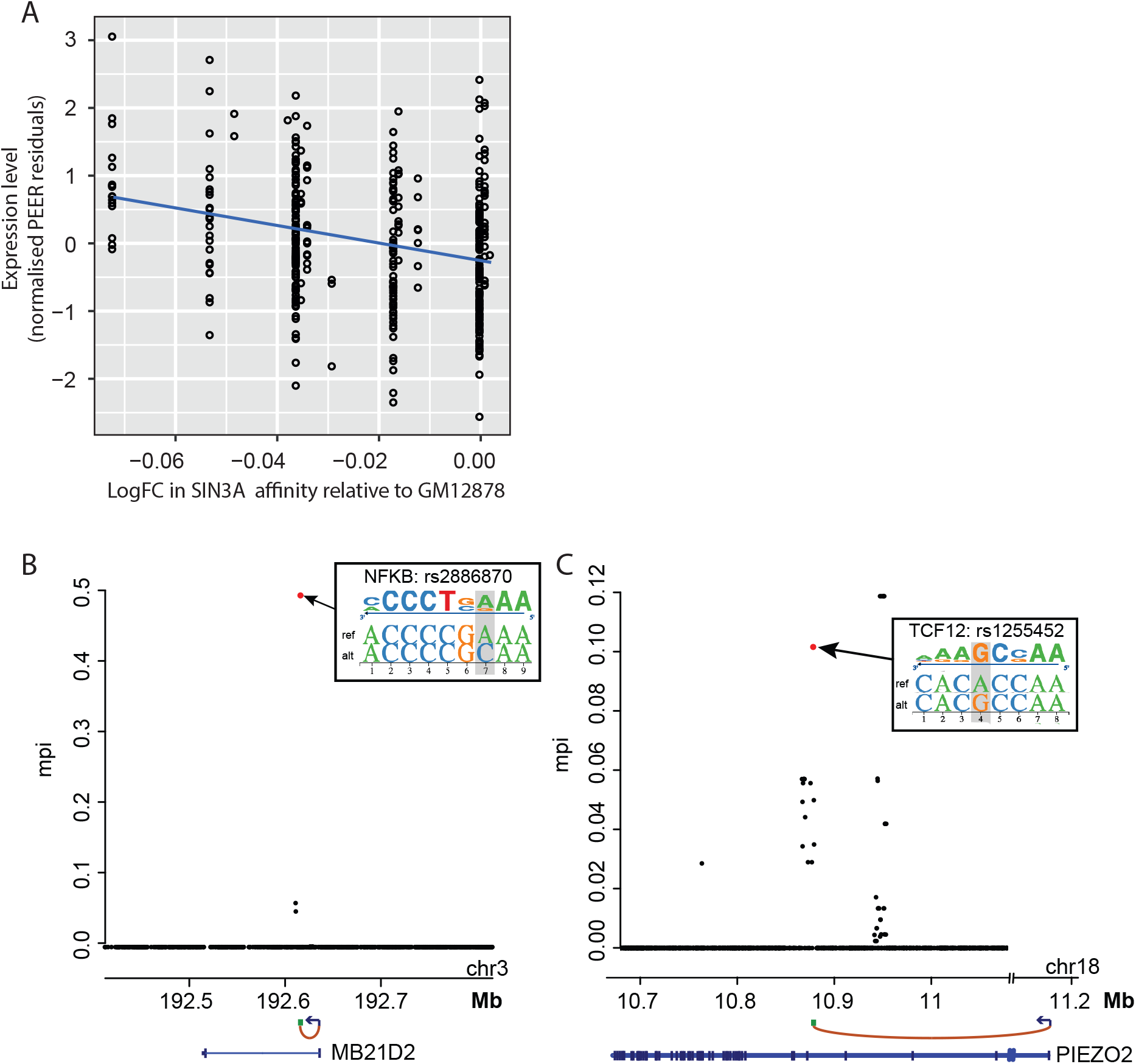
Threshold-free approach for detecting TF binding affinity variant associations with gene expression and validation using GUESSFM. (A) Association between log fold affinity change in CRM affinity for SIN3A relative to the highest affinity allele of GM12878 and mRNA level (normalised PEER residuals) of the connected gene, *SLC23A* (gene level FDR adjusted p-value=1.60×10^−4^, beta=–0.14). (B) Examples of loci, whereby the SNP predicted to have the strongest impact on a CRM’s binding affinity for a given TF has been fine-mapped as a potentially causal variant driving the locus’s association with the expression of a physically connected target gene (GUESSFM marginal posterior probability of inclusion [mppi]>>0.001). Left panel: eGene: *MB21D2;* eQTL rs2886870, predicted to affect NFKB binding affinity. Right panel: eGene: *PIEZO2;* eQTL rs1255452, predicted to affect TCF12 binding affinity. See insets for the effects of the SNPs on the respective TF PWMs.

### TF-binding affinity variants are highly enriched for fine-mapped causal eQTLs

We asked what proportion of TF-binding variants showing association in our analysis could be fine-mapped as causal purely based on the pattern of association signals in their vicinity, without *a priori* prioritisation and pooling of variants per CRM. To this end, we supplied genotype information for +/−200 kb windows around the CRMs with detected associations and the respective gene expression data to GUESSFM, a Bayesian fine-mapping approach that accounts for possible multiple causal variants per locus (74). GUESSFM identified at least one causal variant in ~38% of the analysed CRMs (1807/4718); associations in the remaining CRMs likely could not be fine-mapped due to a lack of statistical power. In ~30% (548/1807) of CRMs with successful fine-mapping, the TF binding variant showing the strongest association per CRM was ranked as possibly causal (marginal posterior probability of inclusion [mppi] >> 0.001), and in the majority of such cases (477/548) this variant was also ranked by GUESSFM among the top five highest-scoring variants in the window (Table S3 and Fig 5B and C for examples). In contrast, just 2.6% (48/1807) random variants within the same windows (matched by allele frequency) were detected as potentially causal by GUESSFM, corresponding to a very significant enrichment of fine-mapped variants for those affecting TF binding (Fisher test p=10^−126^).

### Many CRMs associated with distal gene expression show features of epromoters

We noted that a large number of distal CRMs showing association between TF binding affinity and target gene expression (224 CRMs, 243 TF-CRM variants; Table S4) and connecting to the distal gene promoters in 3D based on PCHi-C also mapped in close proximity (within 200 bp) of the TSS of either one or more other genes (165 and 59 CRMs, respectively, and 284 eGenes; note that the number of eGenes is greater than that of CRMs due to some CRMs mapping in close proximity of multiple TSSs). The absolute majority (87%) of these CRMs localised within chromatin segments with the characteristic features of gene promoters (Figure 6A). Taken together, this suggested that some promoter regions might act as distal regulatory regions of other genes, whose promoters they physically contact. This class of CRMs with dual promoter and activity were independently identified in two recent studies (63, 64). We shall follow Dao et al. (63) in referring to these CRMs as ‘epromoters’.

**Figure 6.**
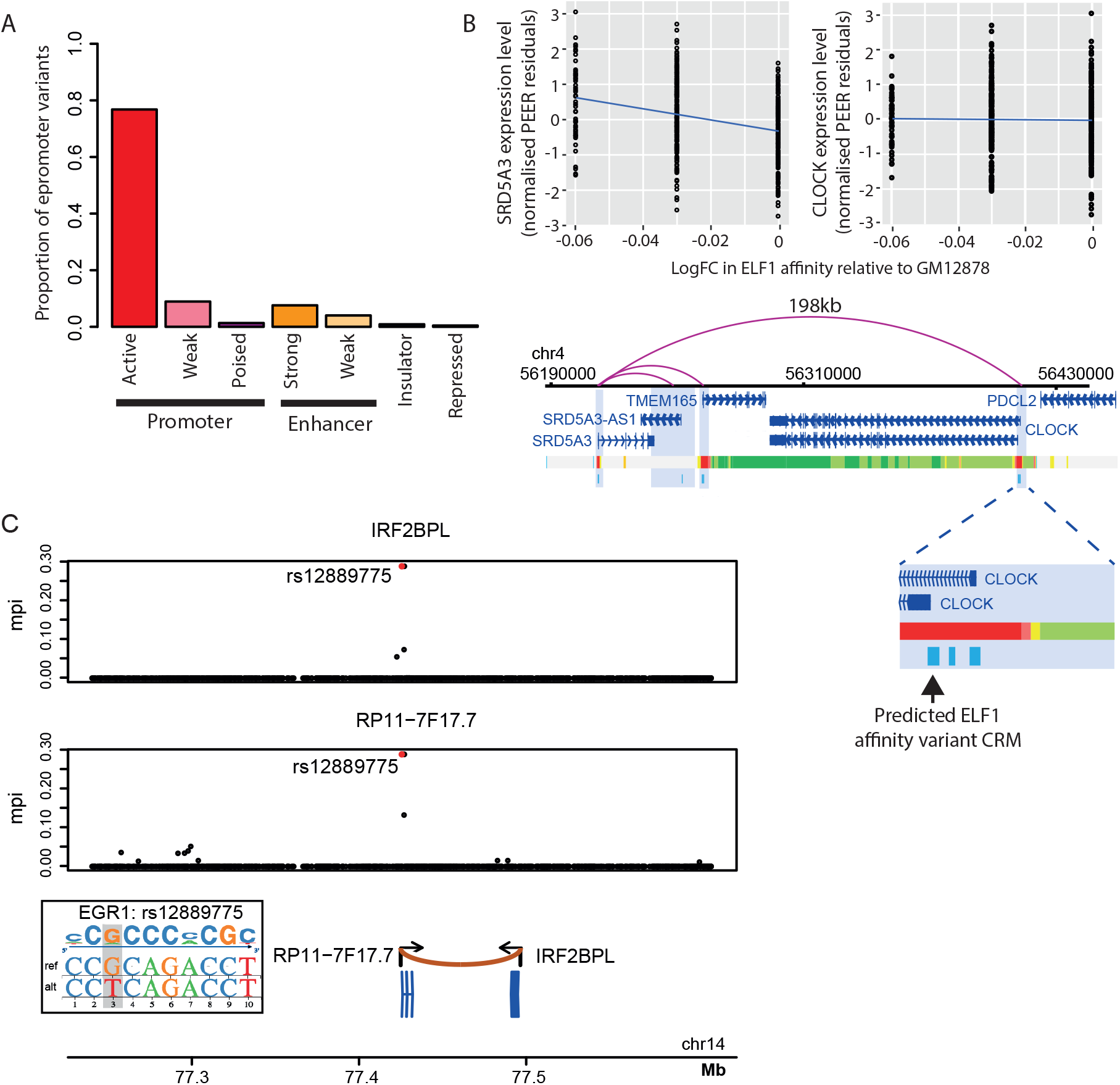
TF-binding affinity variants highlight transcriptional regulatory effects of epromoters. (A) Barplot showing the proportion of distal CRMs showing association between TF binding affinity and target gene expression that map in close proximity (within 200 bp) of another genes TSS overlapping each genome segmentation category (108) for GM12878. (B) Genome browser representation of the distal interactions detected by promoter capture Hi-C (49) for SRD5A3, with CRMs identified at each fragment as well as the proximal window depicted in light blue. The genome segmentation track for GM12878 based on chromHMM (108) is also shown. Enlarged view of an interacting fragment containing three CRMs, one of which harbours variants predicted to impact ELF1 binding affinity and overlaps with the promoter of CLOCK. (B) The association between logFC in CRM affinity for ELF1 relative to the highest affinity allele of GM12878 and mRNA level (normalised PEER residuals) of SRD5A3 and CLOCK. (C) Colocalisation analysis showing shared association between epromoter-located SNP rs12889775 and the expression of both its distal and proximal genes (*IRF2BPL*, top; and lncRNA *RP11−7F17.7*, bottom, respectively); Posterior probability of shared association estimated by the coloc software p_H4_=0.997. This SNP is predicted to affect the epromoter’s binding affinity for EGR1 (see inset).

Most genes located in the immediate vicinity of the identified epromoters were appreciably expressed in LCLs (232/284, 82%). However, TF-binding variation at nearly two-thirds of epromoters whose proximal gene was expressed (139 variants, 64.7%; see Table S2) showed detectable association with a distal gene alone in independent tests (assessed with the threshold-free approach). For example, variation in ELF1 binding affinity at a CRM that shows promoter-associated chromatin marks and localises within 200 bp from the TSS of *CLOCK* gene does not affect *CLOCK* expression. Instead, it associates with expression of *SRD5A3* located 198 kb away, whose promoter it contacts in 3D as detected by PCHi-C (Figure 6B; *SRD5A3:* gene-level FDR = 3.33×10^−21^, ELF1 elastic net p-value=0, ELF1 elastic net beta = −0.21; *CLOCK:* gene-level FDR = 0.88).

The remaining 76 TF-epromoter CRM variants showed associations between with the expression levels of both distal and proximal genes. To obtain formal evidence that these associations were indeed driven by the same variant and not by different variants in LD with each other, we used colocalisation analysis (75), while accounting for multiple independent associations (see Materials and Methods). We submitted to this analysis the most tractable subset of 7 epromoters, for which the association of the respective TF-binding variant with distal gene expression was independently confirmed by fine-mapping (GUESSFM mppi>0.001). At 6/7 analysed epromoters, we found prevailing evidence of shared association signals for both proximal and distal gene (p_H4_>0.66; Table S5). An example of such high-confidence shared signal is variation in EGR1 binding affinity in the epromoter of lncRNA *RP11-71F7.7* that associates with the expression of both *RP11-71F7.7* and another gene, *IRF2BPL* (Fig. 6C). The promoters of these two genes, transcribed in a convergent orientation, are approximately 69 kb apart and contact each other in 3D as detected by PCHi-C.

Taken together, our findings confirm long-range transcriptional regulation by epromoters and suggest that regulatory variants within these elements may have both shared and independent effects on the expression of their proximal and distal target genes.

## Discussion

In this study we have generated an atlas of CRM variants predicted to affect TF binding in LCLs and established their associations with the expression of their target genes identified on the basis of 3D chromosomal interactions or immediate spatial proximity. Notably, we found that many TF binding variants showing associations with distal gene expression localise to the promoters of other genes, providing additional support for the recently characterised class of “epromoter” regulatory elements (63, 64).

We previously reported a collection of TF binding variants for ENCODE-profiled TFs based on 1000 Pilot data (179 individuals) (32). The atlas of binding variants generated in this study is based on a more than two-fold larger sample of EUR individuals from 1000 Genomes release. Importantly, in this study we also used a biophysical model (39) that aggregates TF binding affinities across the whole CRM to increase sensitivity. This in contrast to our previous work (32) and other published resources such as Haploreg (77) and SNP2TFBS (78) that is based on detecting individual PWM motif matches within each region of interest. Our current approach, however, still relies on PWMs to estimate affinities at each sequence window. While this is a straightforward strategy that produces readily interpretable results, it has several limitations. First, conventional PWMs may not adequately describe the binding preferences of some TFs, leading to either overfitted or overly degenerate models. For such TFs, k-mer-based approaches to modelling sequence consensus (79) or models incorporating dependencies between motif positions (80–82) may be more appropriate. In addition, it was recently shown that modelling DNA shape may aid binding prediction for some TFs with seemingly degenerate sequence preferences (83). Moreover, even for TFs whose binding can be modelled by PWMs reasonably well, variants affecting their binding in vivo may be based outside of the immediate binding consensus, possibly due to cooperative effects (25, 26, 30). At the expense of interpretability, this problem can be mitigated by using machine learning models such as DeFine (84) and DeepSEA (85), with the latter approach employed in a recent study investigating impact of regulatory variants on gene expression (86). Alternatively, effects on TF binding can be obtained directly from allele-specific or multi-individual TF binding data (25–28, 30, 87).

To identify the putative target genes of remote regulatory variants, we have capitalised on Promoter Capture Hi-C (PCHi-C) - a high-resolution technique for global chromosomal interaction profiling (48–50). Our approach is based on the classic model of long-range transcriptional control that necessitates physical contacts between enhancer elements and their target promoters through DNA looping (44, 45), which has been validated by a number of approaches, most recently by in vivo imaging (88–90). However, recent evidence suggests that at least some regulatory regions may exert action on their target promoters without coming into proximity with them (91, 92). Alternative methods of assigning the target genes of regulatory regions, such as those based on correlated chromatin activity of promoters and enhancers (20, 93–96) may account for the effects of these regions. Emerging high-throughput functional screens of cis-regulatory activity (8, 9, 64, 97) are proving to be instrumental for direct functional validation of enhancer targets identified with either approach.

In our study, we restricted the analysis to CRMs with predicted variation in TF binding and their putative proximal and distal target genes, which has considerably reduced multiple testing burden and made it possible to combine the effects of multiple polymorphisms within the same CRM based on TF binding models. This analysis framework has yielded highly sensitive and interpretable associations for pre-selected loci. However, owing to its selective nature, it is by no means a substitute to conventional association testing. Theoretically, the effects of nucleotide variants on TF binding can also be incorporated as a prior in global association analyses such as fgwas (98), and have already been used in eQTL fine-mapping (99). We have previously confirmed that promoter-interacting regions are strongly enriched for eQTLs of the physically connected distal genes (51). An optimal statistical framework for incorporating 3D interaction data into eQTL testing is, however, yet to be established.

Our finding that polymorphic TFBSs at distal CRMs show gene expression associations less frequently compared with proximal regions is consistent with the high degree of redundancy of long-range regulatory elements (5–7, 100, 101). Predicting the extent of buffering of regulatory variation for a given CRM with a reasonable precision is an important problem that is currently highly challenging due to the sheer number of parameters and the relatively small sample sizes of multi-individual expression datasets. Profiling gene expression in the emerging much larger genotype panels such as UK10K (102) may provide opportunities for addressing this question.

We observed that a large proportion of CRMs showing associations with the expression of physically connected distal genes localised in the promoter regions of other genes. This provides additional evidence to the recently characterised class of “epromoters”: elements with a dual proximal and distal activity that were discovered on the large scale using high-throughput reporter and CRISPR knockout screens (63–65). Empirically, chromosomal interactions between epromoter CRMs and their distal targets fall into the category of promoter-promoter interactions. Until recently, these interactions have been viewed primarily in the context of coordinated gene activation or repression (103–105), such as that observed in Hox and histone clusters (103, 106). That some promoter-promoter contacts reflect epromoter-distal target gene relationships suggests that these contacts may show functionally and possibly even structurally distinct properties.

We show that TF-binding variation at epromoters may or may not co-associate with the expression of both proximal and distal genes at the same time. Shared association is consistent with the findings from massively parallel reporter assays that the same sequences are often involved in mediating both promoter and enhancer activity in vitro (107). It is possible that some non-shared effects observed in our study in vivo are underpinned by the role of the affected TFs in mediating long-range contacts. Additionally, epromoter elements may show different degrees of redundancy with respect to the proximal and distal target gene.

Overall, our analysis demonstrates the potential of model-based prioritisation and pooling of variants a priori of testing for increasing the sensitivity of identifying individual associations and revealing their shared biological properties.

## Data Availability

The list of the detected TF affinity CRM variants, the full data on CRM variant – gene expression associations and the raw output of GUESSFM fine-mapping have been uploaded to Open Science Framework (https://osf.io/fa4u7/).

## Supplementary Data

**Figure S1.** Suitability of the association signal colocalisation algorithm for analysing pairs of signals within the same dataset.

**Table S1.** Significant associations between TF-binding affinity CRM variants and target gene expression identified with the thresholded approach.

**Table S2.** Significant associations between TF-binding affinity CRM variants and target gene expression identified with the threshold-free approach.

**Table S3.** Posterior probabilities of the prioritised TF-binding affinity eQTLs being causal estimated by GUESSFM fine-mapping algorithm.

**Table S4.** A list of epromoters with detected TF-binding affinity variation.

**Table S5.** Association signal colocalisation analysis for the proximal and distal target genes of selected epromoters.

## Acknowledgements

The authors wish to thank Paula Freire-Pritchett, Hashem Koohy, Jonathan Cairns and Simon Andrews for advice and technical assistance, and all members of MS lab for helpful discussions.

