## Supplementary material for "Functional effects of variation in transcription factor binding highlight long-range gene regulation by epromoters": Figure S1

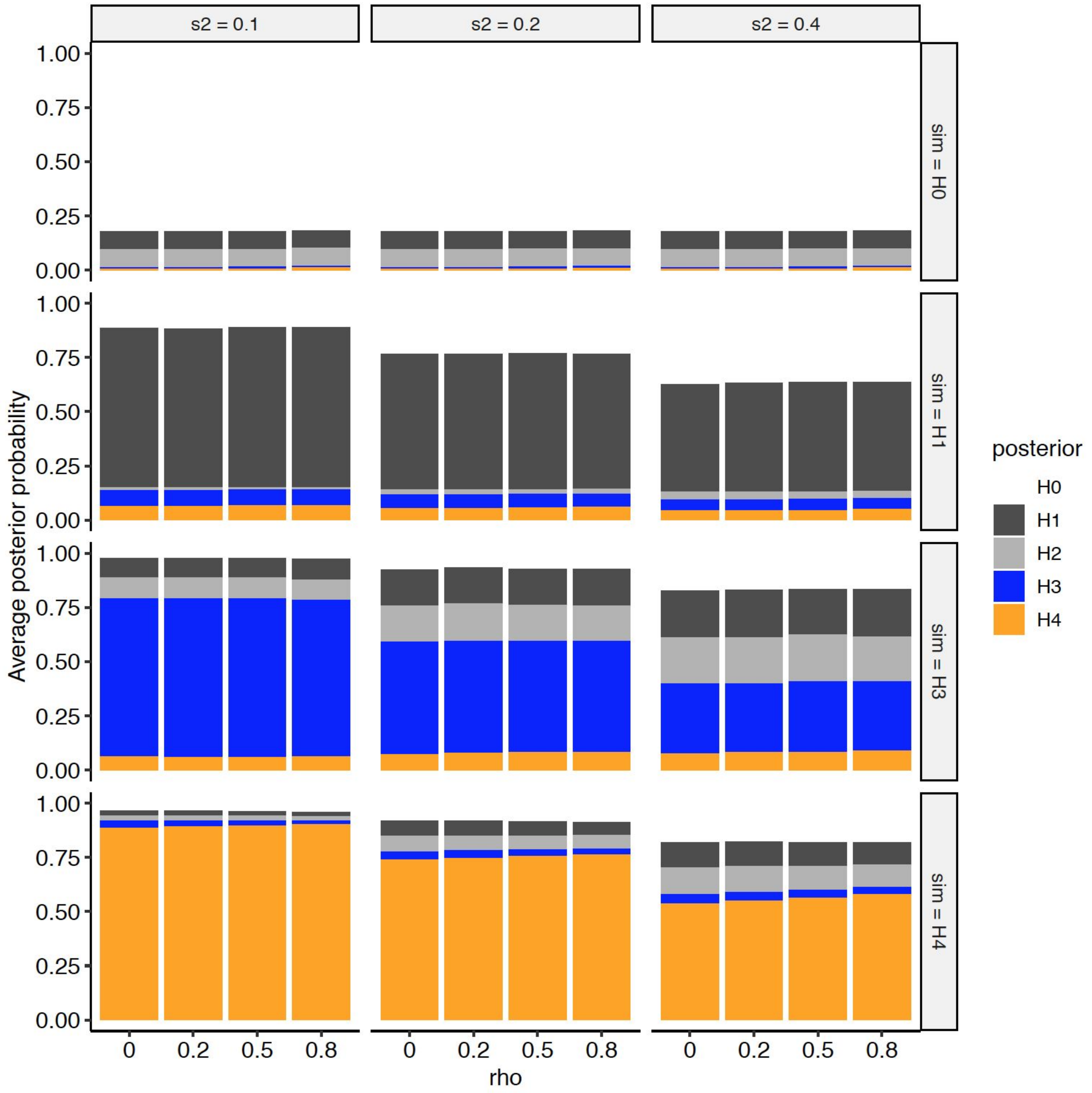


**Figure S1. Suitability of the association signal colocalisation algorithm for analysing pairs of signals within the same dataset.** We sampled 200 individuals from 1000 Genomes and simulated two quantitative traits with varying levels of non-genetic trait variance (s2, with higher s2 implying lower power) and residual correlation after main genetic effects were accounted for (rho). Genetic effects were simulated as either sim=H0 (no effects for either trait), sim=H1 (single causal variant for trait 1 only), sim=H3 (different single causal variant for each trait), sim=H4 (the same single causal variant for each trait). Barplots show the average posterior probability of each of the five possible hypotheses, and show that increasing rho has a negligible impact on inference, reassuring that it is appropriate to use coloc to compare expression of two genes quantified on the same individuals.
